# High-resolution map of the Fc-functions mediated by COVID-19 neutralizing antibodies

**DOI:** 10.1101/2023.07.10.548360

**Authors:** Ida Paciello, Giuseppe Maccari, Elisa Pantano, Emanuele Andreano, Rino Rappuoli

**Affiliations:** Monoclonal Antibody Discovery (MAD) Lab, Fondazione Toscana Life Sciences, Siena, Italy; Data Science for Health (DaScH) Lab, Fondazione Toscana Life Sciences, Siena, Italy; Department of Biotechnology, Chemistry and Pharmacy, University of Siena, Siena, Italy; Fondazione Biotecnopolo di Siena, Siena, Italy

## Abstract

A growing body of evidence shows that Fc-dependent antibody effector functions play an important role in protection from severe acute respiratory syndrome coronavirus 2 (SARS-CoV-2) infection. To unravel the mechanisms that drive these responses, we analyzed the phagocytosis and complement deposition mediated by a panel of 482 human monoclonal antibodies (nAbs) neutralizing the original Wuhan virus, expressed as recombinant IgG1. Our study confirmed that nAbs no longer neutralizing SARS-CoV-2 Omicron variants can retain their Fc-functions. Surprisingly, we found that nAbs with the most potent Fc-function recognize the N- terminal domain, followed by those targeting Class 3 epitopes in the receptor binding domain. Interestingly, nAbs direct against the Class 1/2 epitopes in the receptor binding motif, which are the most potent in neutralizing the virus, were the weakest in Fc-functions. The divergent properties of the neutralizing and Fc- function mediating antibodies were confirmed by the use of different B cell germlines and by the observation that Fc-functions of polyclonal sera differ from the profile observed with nAbs, suggesting that not-neutralizing antibodies also contribute to Fc-functions. These data provide a high-resolution picture of the Fc-antibody response to SARS-CoV-2 and suggest that the Fc contribution should be considered for the design of improved vaccines, the selection of therapeutic antibodies and the evaluation of correlates of protection.

## INTRODUCTION

The coronavirus diseases 2019 (COVID-19) pandemic has been responsible for more than 768 million infections and nearly 7 million deaths reported worldwide^1^. Significant progress in the fight against SARS-CoV-2 has been achieved with the approval and the administration of vaccines and monoclonal antibodies (mAbs). Antibodies administrated or elicited by vaccination neutralize viral entry into host cells by the interaction between the antigen-binding fragment (Fab) region of the antibodies and the SARS-CoV-2 Spike (S) protein^2, 3^. Early predictive models suggested that neutralizing levels of anti-S protein antibodies elicited by vaccination correlate to protection from infection with SARS-CoV-2^4^. These studies were conducted when the S protein antigen encoded by COVID-19 vaccines was the same as the circulating virus. Unfortunately, during the COVID- 19 pandemic, SARS-CoV-2 variants developed progressively several mutations mainly placed in the receptor binding domain (RBD) and the N-terminal domain (NTD) of the S protein^5^. These SARS-CoV-2 variants, and especially the Omicron lineages, show resistance against the majority of monoclonal antibodies and the ability to evade infection and vaccination-induced immunity^6, 7^. As a result, higher neutralizing antibody titers are necessary to induce protection from SARS-CoV-2 variants^8^. Despite elevated resistance to neutralization, individuals infected with recently emerged variants showed lower severity of disease, suggesting that additional components of the immune system play a role in protection from severe COVID-19^9^. Fragment crystallizable (Fc)-dependent antibodies effector functions are emerging as key players in the determining the outcome of severe infection^10, 11^. Several evidences report that the Fc-functions provide protection from COVID- 19 diseases even in the absence of neutralization^10–12^. Indeed, Addetia et al demonstrated that the FDA approved mAb S309 (sotrovimab) triggers antibody-dependent cell cytotoxicity (ADCC) in vitro and protects mice against BQ.1.1 challenge in vivo despite loss of neutralization activity^12^. In addition, Mackin et al reported that mice lacking expression of activating FcγRs, especially murine FcγR III (CD16), or depleted of alveolar macrophages, lose the antiviral activity of passively transferred immune serum against multiple SARS-CoV-2 variants^13^. Recent serology data have also demonstrated that SARS-CoV-2 vaccine induced higher Fc-receptor binding antibody levels and consequently higher humoral and cellular immune responses in subjects with previous SARS-CoV-2 infection compared to infection of naïve individuals. In addition to SARS-CoV-2, several studies reported that Fc-effector functions are also crucial to confer protection against other pathogens including Influenza virus^16, 17^, Ebola virus^18^, as well as several bacterial infections^19^. Given the relevance of the Fc-mediated functions, it becomes of utmost importance to understand which are the epitopes and B cell germlines involved in this response and if they overlap with those inducing neutralizing antibodies. To answer this question, we took advantage of our unique panel of 482 Wuhan neutralizing human monoclonal antibodies (nAbs), most of which lost their neutralization activity against Omicron variants^20^. These antibodies derived from naïve donors who were immunized with two mRNA vaccine doses (Seronegative, SN2) and subsequently re-enrolled after receiving the third vaccine dose (Seronegative, SN3), and convalescent donors who had been infected before vaccination (Seropositive, SP2)^21, 22^. Antibody-dependent cellular phagocytosis (ADCP) and complement-dependent deposition (ADCD) were used to characterize at a single-cell level the Fc-effector functions of our nAb panel. These functions depend on antibody isotype, location and geometry of the antibody-binding epitope, stoichiometry of the antibody to antigen and affinity of the antibody for both its antigen and its cognate FcR^11^. To investigate the influence of the SARS-CoV-2 S protein domains and epitopes to induce Fc-activities, all nAbs were expressed as recombinant IgG1 isotype. Our data revealed that nAbs that lost their neutralization activity can retain Fc-functions against tested Omicron variants and shared key features among infected and vaccinated donors. In particular, different domains, epitopes and B cell germlines were shown to drive Fc-functions and the neutralization response. These data suggest that synergy between Fc-mediated functions and neutralization could be a successful strategy to design improved vaccines and therapeutics against COVID-19.

## RESULTS

### Hybrid immunity induces high Fc-dependent effector functions polyclonal antibody response

To explore the polyclonal response elicited by infection and vaccination, we collected plasma samples from seronegative subjects immunized with two (SN2, n=5) or three (SN3, n=4) mRNA vaccine doses^21, 22^, and seropositive donors who had been infected before mRNA vaccination (SP2, n=5). As first step, we evaluated the ability of the polyclonal antibody response to drive cellular phagocytosis through an antibody dependent cellular phagocytosis (ADCP) assay, evaluated by measuring the THP-1 engulfment of beads coated with fluorescent S proteins, (**Fig. 1A, Extended Data Figure 1A**). SP2 induced the highest ADCP activity to all SARS- CoV-2 S proteins, followed by SN3 and SN2 (**Fig. 1B**). The ADCP activity slightly decreased with BA.1 and had a dramatic reduction with BA.2. However, the ADCP activity of SP2 remained the highest (**Fig. 1B**). Next, we evaluated the antibody dependent complement deposition (ADCD) response in SN2, SN3 and SP2. As illustrated in Fig. 1C, Expi293F cells expressing SARS-CoV-2 Wuhan, BA.1 or BA.2 S protein were incubated with plasma samples and the C3 deposition was measured through flow-cytometry (**Extended Data Figure 1B**). In line with ADCP results, SP2 subjects showed the highest ADCD activity followed by SN3 and SN2 (**Fig. 1D**). The Omicron variants showed a drastic reduction in ADCD activity in all cohorts with almost complete evasion of this response in SN2 and SN3. In fact, BA.1 showed a 2.5-fold, 3.2-fold and 4.1-fold ADCD reduction in SN2, SN3 and SP2 respectively. Similarly, the use of BA.2 S protein led to 2.1-fold, 2.7-fold and 3.3-fold ADCD decrease in SN2, SN3 and SP2 respectively (**Fig. 1D**).

**Figure 1.**
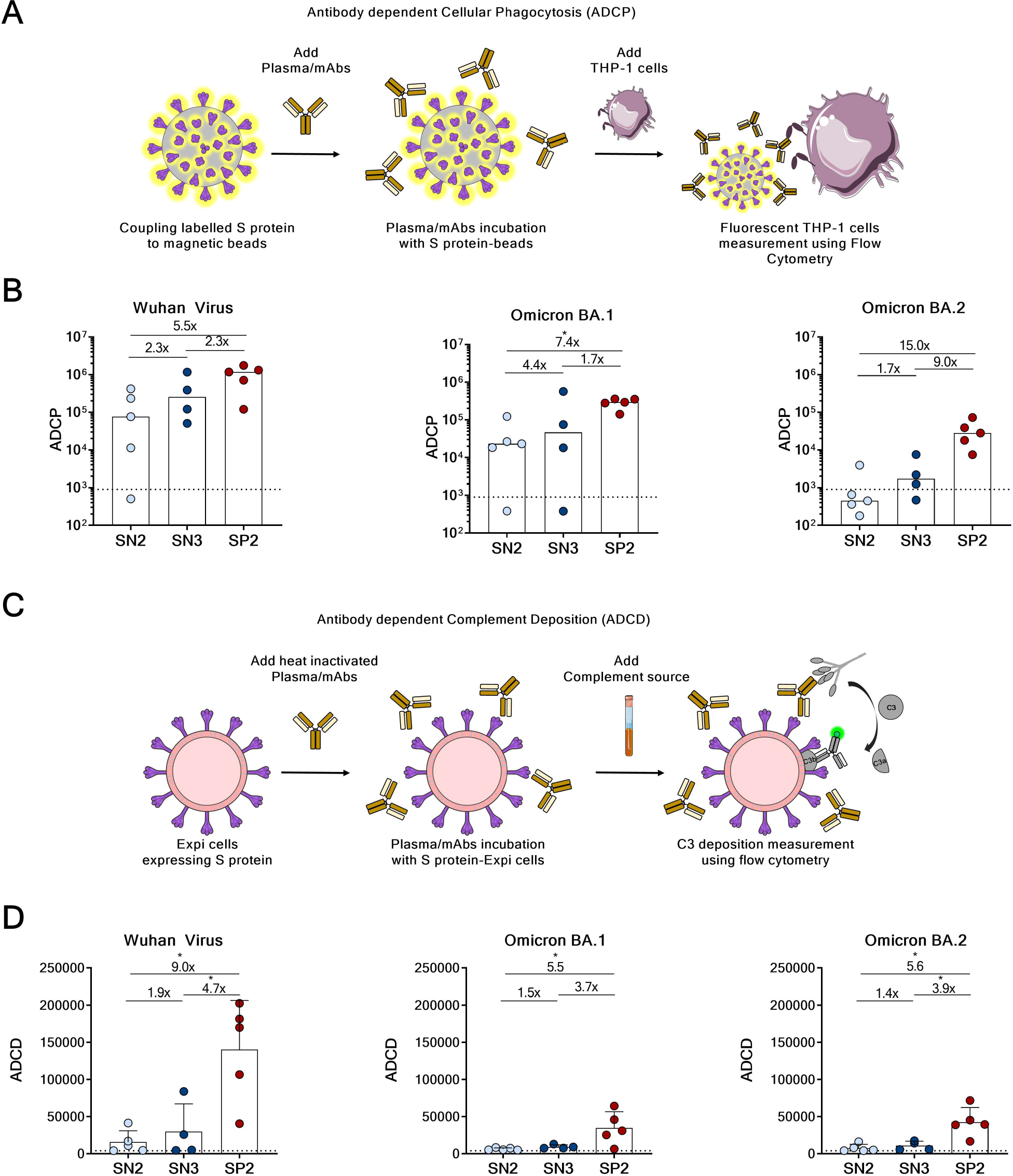
Antibody Dependent Cellular Phagocytosis (ADCP) and Antibody Dependent Complement Deposition (ADCD) driven by polyclonal response. (A) Schematic representation of the ADCP assay. (B) Comparison of ADCP activity against SARS-CoV-2 WT, BA.1 and BA.2 variants in THP-1 incubated with plasma derived from SN2, SN3 and SP2 cohorts. (C) Illustration of the ADCD assay performed. (D) Comparison of ADCD activity of plasma derived from SN2, SN3 and SP2 cohorts against SARS-CoV-2 WT, BA.1 and BA.2 variants. Threshold of positivity has been set based on the value of the negative control (dotted line) and non- parametric Mann–Whitney t-test was used to evaluate the statistical significance between groups. Two-tailed p value significances are shown as *p<0.05.

### ADCP is driven by NTD-targeting nAbs after a third mRNA vaccine dose and hybrid immunity

Next, we characterized the ADCP of neutralizing monoclonal antibodies (nAbs) isolated from SARS-CoV-2 naïve donors after two (SN2; n=52), three vaccinations (SN3, n=206) and 224 nAbs derived from subjects vaccinated after COVID-19 infection (SP2)^21, 22^. Therefore, our total nAb panel is composed by 482 antibodies neutralizing the Wuhan virus of which 74-98% had lost the ability to neutralize Omicron BA.1 and BA.2 variants^20^. All these nAbs were expressed as IgG1. Therefore, the Fc-functions observed depend on the antibody-binding epitope site, the stoichiometry and affinity of the antibody Fab region for its antigen. Up to 99.4% (n=479) of nAbs induced phagocytosis activity against the Wuhan SARS-CoV-2 S protein (**Fig. 2A**). However, the overall potency of ADCP in SN2 are significantly lower than that of SN3 and SP2 (SN2 vs SN3 p<0.0001; SN2 vs SP2 p<0.001) (**Fig. 2A**). Over 70 and 60% of nAbs in SP2 and SN3 cohorts showed high ADCP activity respectively, while only 35% of this class of antibodies was found in the SN2 group. For this screening, an unrelated plasma was used as negative control and the threshold for sample positivity was set at 3-fold the ADCP score of the negative control. Based on this cutoff point, nAbs were classified as high (>8-fold the threshold), medium (> 4-fold the threshold), or low (up to 4-fold the threshold) phagocytosis inducers and given the extend of our nAb panel, antibodies were tested at a single-point dilution. To confirm that ADCP titers obtained were independent from the antibody concentration, a nAb dose-phagocytosis titer correlation was performed. The analyses showed that the S protein-dependent opsonization levels were unrelated to the antibody concentration used (**Extended Data Fig. 2A**). Moreover, additional correlation analysis revealed that there was no correlation between the neutralization and ADCP titers (**Extended Data Figure 2B**). Then, we tested the ADCP against the Omicron BA.1 and BA.2 lineages. Overall, nAbs tested displayed a significantly lower ADCP activity against the Omicron variants compared to Wuhan (**Fig. 2A**). Only 15.5 (9/52) and 25.0% (56/224) of nAbs in the SN2 and SP2 cohort respectively retained ADCP against BA.1. Conversely, up to 60.7% (125/206) of tested antibodies in the SN3 group drove phagocytosis although with much lower ADCP titers. A different scenario was observed against the Omicron BA.2, where only 25.7% (53/206) of nAbs derived from the SN3 cohort induced opsonization which was similar to what observed in the SN2 group (7/52; 13.5%). An opposite trend was observed in the SP2 cohort, where 31.7% (71/224) of nAbs drove ADCP against the Omicron BA.2 which was significantly higher to what observed in the SN3 group. In our previous work, we identified the S protein domain recognized by our nAb panel. To better understand the impact of S protein domains, we analyzed the ADCP activity of the 369 RBD-binding antibodies (SN2=37, SN3=154, SP2=178), the 89 NTD-targeting nAbs (SN2=9, SN3=43, SP2=37) and the 24 antibodies that bind to the S protein trimer (SN2=6, SN3=9, SP2=9)^20^. We exploited this information to understand which domains were mainly responsible for ADCP. Our data showed that almost all nAbs, independently from the targeted domain, were able to induce phagocytosis against the Wuhan SARS-CoV-2 virus. However, both ADCP frequency (**Fig. 2B-D, left panel**) and potency (**Fig. 2B-D, right panel; Extended data Table 1**) are drastically decreased versus the Omicron BA.1 and BA.2. In the SN2 cohort around 20% of nAbs showed to induce phagocytosis against tested variants (**Fig. 2B, left panel**). SN3 and SP2 showed higher ability to retain ADCP against BA.1 and BA.2 even if different trends were observed. Indeed, SN3 drove higher phagocytosis to BA.1 compared to BA.2, with NTD nAbs being the most endurable (79%), followed by S protein (66.9%) and RBD (55.2%) binders (**Fig. 2C, left panel**). As for SP2, higher phagocytosis to BA.2 compared to BA.1 was observed. S protein targeting nAbs were the most resistant to BA.2 (44.4%) followed by NTD (37.6%) and RBD (29.8%) binding antibodies (**Fig. 2D, left panel**). Following, we evaluated the ADCP potency of our nAb panel (**Fig. 2B-D, right panel**). Overall, low phagocytic activity was observed in the SN2 group confirming that third vaccination is crucial to increase ADCP potency (**Fig. 2B-C, right panel**). In the SN3 cohort, NTD-targeting nAbs showed the highest ADCP against Wuhan and BA.1, while similar potency was observed to BA.2 when compared to S protein and RBD binding nAbs (**Fig. 2C, right panel**). Similarly, NTD- targeting nAbs derived from SP2 donors showed a significantly higher ADCP potency against Wuhan and BA.2 compared to S protein and RBD antibodies, while similar ADCP activity was observed against BA.1 (**Fig. 2D, right panel**).

**Figure 2.**
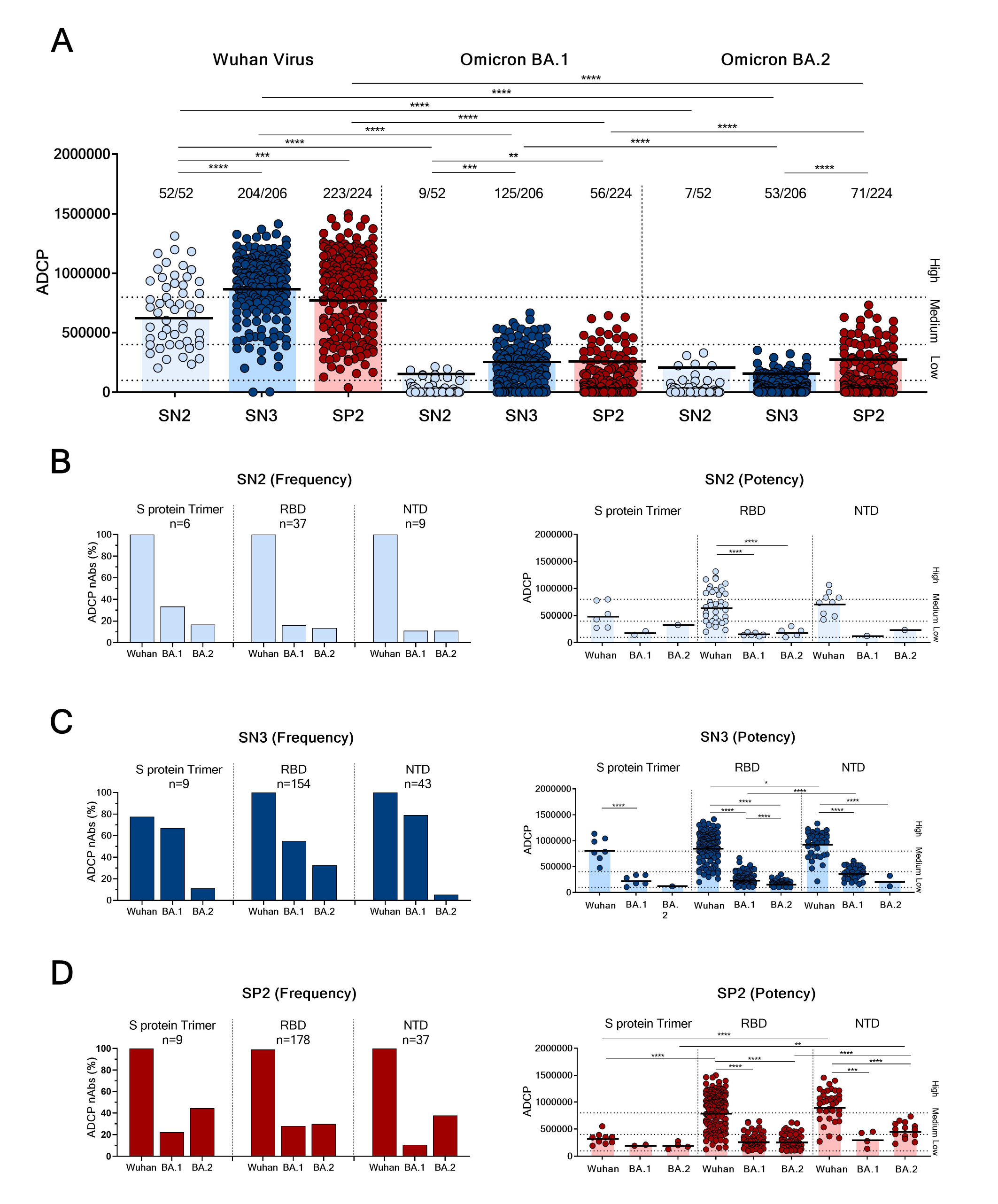
ADCP driven by neutralizing monoclonal antibodies. (A) Phagocytosis induced by neutralizing antibodies isolated from SN2, SN3 and SP2 cohorts against the original SARS-CoV-2 original Wuhan virus, BA.1 and BA.2 was described. (B-D) Bar graphs show the distribution of ADCP frequency between RBD, NTD domains and the S protein in trimeric conformation against SARS-CoV-2 original Wuhan virus, the Omicron BA.1 and BA.2 variants in SN2 (B, left panel), SN3 (C, left panel) and SP2 (D, left panel) cohorts. Dot plots show the ADCP potency comparison of S protein trimer, RBD and NTD binders against SARS-CoV-2 VoCs in SN2 (B, right panel), SN3 (C, right panel) and SP2 (D, right panel) cohorts. Non-parametric Mann–Whitney t-test was used to evaluate the statistical significance between groups. Two-tailed p value significances are shown as *p<0.05, **p < 0.01, ***p < 0.001, and ****p<0.0001.

### NTD-targeting antibodies are major ADCD drivers after a third mRNA vaccine dose

As next step, we measured the SARS-CoV-2 nAbs ability to stimulate complement deposition. The activation of complement on SARS-CoV-2 antigen-antibody complex was measured using S protein expressed on the surface of target cells and detecting C3 complement deposition in presence of baby rabbit complement source. The same nAb panel described above was used to evaluate ADCD. As previously described, based on the mean fluorescent intensity (MFI) value of the negative control, nAbs were classified as high (>8-fold the threshold), medium (>4-fold the threshold), or low (up to 4-fold the threshold) ADCD inducers. Overall, we observed lower ADCD activity in all groups compared to ADCP (**Fig. 2A and Fig. 3A**). SN3 showed the highest ADCD against SARS-CoV-2 Wuhan with up to 64.6% (133/206) of nAbs activity. Differently, only 30.8 (16/52) and 48.7% (109/224) of nAbs from SN2 and SP2 respectively showed ADCD against this virus (**Fig. 3A**). In addition, these two latter cohorts lose almost completely their ADCD activity against the Omicron variants. Only 3.8 (2/52) and 5.8% (3/52) of nAbs in the SN2 cohort showed ADCD against BA.1 and BA.2 respectively, while 7.1 (16/224) and 21.4% (48/224) of nAbs in the SP2 group induced complement deposition against these variants. As for ADCP, we evaluated a possible correlation between ADCD titers, antibody concentration and neutralization titers. Also in this case, we observed that ADCD levels were concentration-independent (**Extended Data Fig. 2C**) and no correlation was observed between neutralization and ADCD titers (**Extended Data Fig. 2D**). Following, we evaluated ADCD induced activity for S protein trimer, RBD and NTD targeting nAbs. NTD binding antibodies were the most active in all three cohorts against the Wuhan virus with 100% of this antibody showing activity in the SN3 cohort (**Fig. 3B-D, left panel**). Compared to ADCP, a more severe loss of ADCD was observed against BA.1 and BA.2 especially in the SN2 and SP2 cohorts while more than 60% of NTD-binders in SN3 maintained their functionality against these variants (**Fig. 3B-D, left panel; Extended data Table 2**). As next step, we evaluated the ADCD potency induced by nAbs isolated from the three cohorts enrolled in this study. In line with previous results, NTD-targeting nAbs derived from SN3 cohort triggered the highest ADCD titers against the original SARS-CoV-2 virus and the Omicron variant BA.1 (**Fig. 3B-D, right panel**). In the SP2 cohort, RBD and NTD Abs binders show similar potency against the Wuhan virus but, the NTD binders were the most powerful nAbs identified against the Omicron variant BA.2 (**Fig. 3D, right panel**).

**Figure 3.**
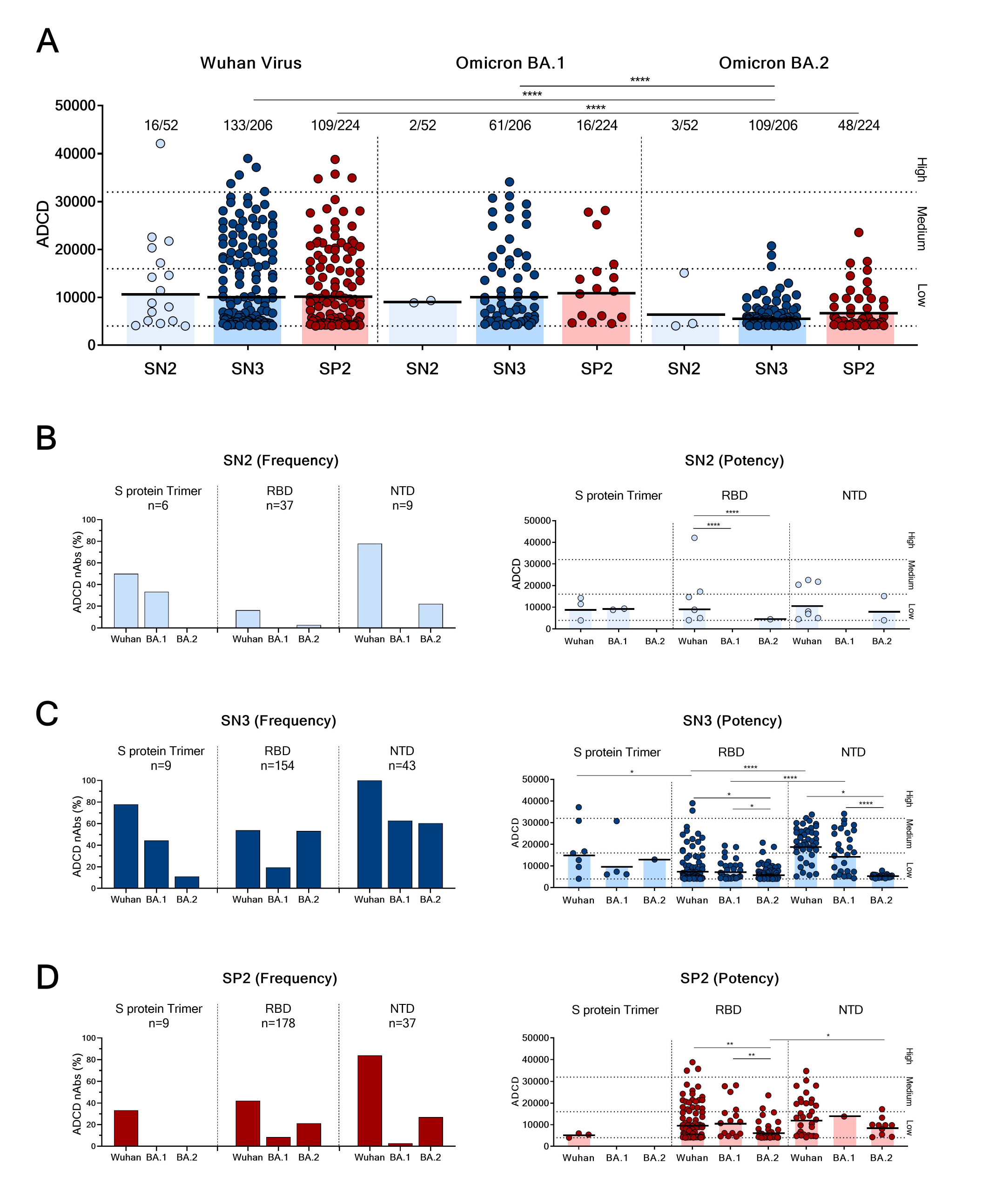
Antibody-dependent complement deposition (ADCD) response mediated by neutralizing monoclonal antibodies. (A) Antibodies dependent complement deposition against the original SARS-CoV-2 original Wuhan virus, BA.1 and BA.2 variants induced by nAbs isolated from SN2, SN3 and SP2 cohorts was described. (B-D) Bar graphs show the percentage of S protein, RBD and NTD nAbs binders able to induce ADCD against SARS-CoV-2 original Wuhan virus, the Omicron BA.1 and BA.2 in SN2 (B, left panel), SN3 (C, left panel) and SP2 (D, left panel) cohort. Dot plots show the ADCD potency of S protein trimer, RBD and NTD antibodies binders against SARS-CoV-2 VoCs in SN2 (B, right panel), SN3 (C, right panel) and SP2 (D, right panel) cohort. Non-parametric Mann–Whitney t-test was used to evaluate the statistical significance between groups. Two- tailed p value significances are shown as *p<0.05, **p < 0.01, and ****p<0.0001.

### RBD Class 3 targeting antibodies induce a strong and variant resistant Fc-mediated response

In our previous study, RBD-targeting nAbs were classified based on their capacity to compete with the Class 1/2 J08^23^, Class 3 S309^24^ and Class 4 CR3022^25^ antibodies (**Fig. 4A**). This information was used to understand the S protein RBD regions that mainly drive the antibody Fc-mediated response. Class 1/2, Class 3 and Class 4 nAbs were respectively n=17, 24 and 0 for SN2, n=98, 30 and 3 for SN3, and n=114, 48 and 7 for SP2^20^. Our results showed that almost all nAbs in the three cohorts induce ADCP against the SARS-CoV-2 Wuhan virus (**Fig. 4B-D**) but these mAbs showed a major drop in ADCP activity against Omicron BA.1 and BA.2. Moreover, Class 3 antibodies show slightly higher potency and broader protection compared to Class 1/2 nAbs especially in SP2 cohort (**Fig. 4B-D**). Indeed, 25.0 and 25.0% in SN2, 70.0 and 63.3% in SN3, and 54.2 and 50.0% in SP2 Class 3 nAbs maintained ADCP against BA.1 and BA.2 respectively. Despite Class 4 nAbs seems to induce slightly higher ADCP potency, the low number of antibodies makes difficult the comparison with the other classes of nAbs herein described. Next, we evaluated the ADCD activity of nAbs belonging to the three RBD Classes of antibodies. As shown in Fig. 4E-G, lower percentages of RBD targeting nAbs induced ADCD in all three cohorts. Anyway, as described for ADCP, Class 3 nAbs induce by a third booster dose or hybrid immunity showed the strongest and broadest ADCD activity against Omicron BA.1 (SN3=40%; SP2=27.1%) and BA.2 (SN3=70%; SP2=43.8%) (**Fig. 4F-G**). The nAb sample size in the SN2 cohort was not sufficient to perform comparative analyses with the other groups. In addition, despite the low number, Class 4 nAbs also showed ADCD levels similar to Class 3 nAbs in both SN3 and SP2.

**Figure 4.**
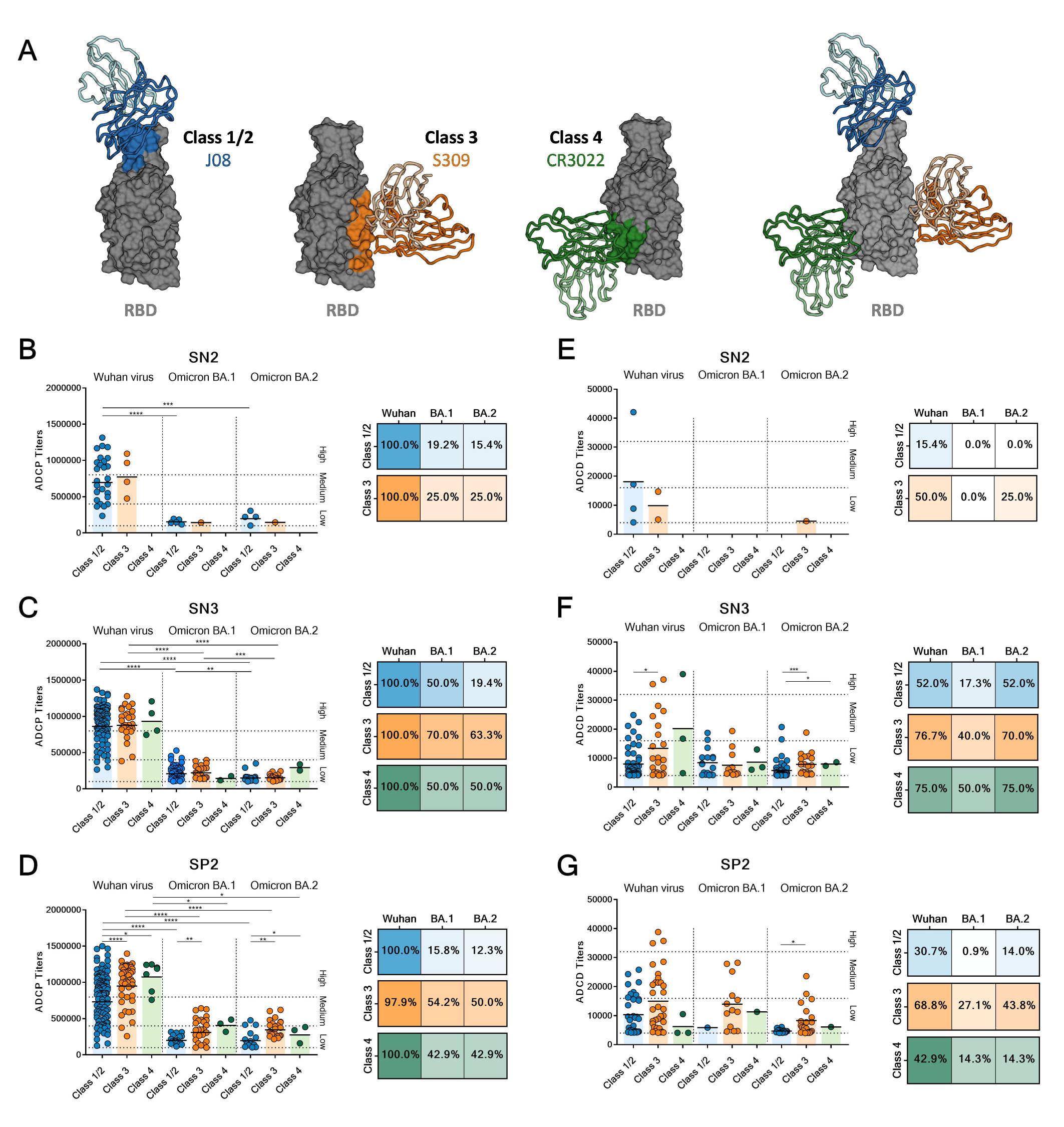
ADCP and ADCD response distribution in RBD-targeting nAbs. (A) Illustration of the epitope regions recognized by Class 1/2 (blue), Class 3 (orange) and Class 4 (dark green) RBD-binding antibodies. (B-D) Dot plots show the titers of nAbs mediated phagocytosis based on their ability to bind Class 1/2, Class 3 and Class 4 regions on the RBD in SN2 (B, left), SN3 (C, left) and SP2 (D, left) cohorts. The heat map next to each dot plot displays the percentage of nAbs able to induce phagocytosis in the SN2 (B, right), SN3 (C, right) and SP2 (D, right) cohorts. (E-F) Dot plots show the ADCD titers of Class 1/2, Class 3 and Class 4 nAbs binders in SN2 (E, left), SN3 (F, left) and SP2 (G, left) cohorts. The heat map linked to each graph display the percentage of nAbs able to induce complement deposition in the SN2 (E, right), SN3 (F, right) and SP2 (G, right) cohorts. Non-parametric Mann–Whitney t-test was used to evaluate the statistical significance between groups. Two-tailed p value significances are shown as *p<0.05, **p < 0.01, ***p < 0.001, and ****p<0.0001.

### Different B cell germlines are responsible for ADCD and ADCP after vaccination and infection

Following the Fc functional characterization, we investigated the predominant B cell germlines and V-J gene rearrangements (IGHV;IGHJ) used by ADCP and ADCD inducing nAbs. From our panel of 482 nAbs, we retrieved 430 heavy chain sequences (SN2=46; SN3=176; SP2=208)^20, 21^. As previously described, the most abundant germlines encoding for neutralizing antibodies were the IGHV1-69;IGHJ4-1, IGHV3-30;IGHJ6-1, IGHV3- 53;IGHJ6-1, IGHV3-66;IGHJ4-1 for SN2, IGHV1-58;IGHJ3-1, IGHV1-69;IGHJ3-1, IGHV1-69;IGHJ4-1, IGHV3-66;IGHJ4-1, and IGHV3-66;IGHJ6-1 for SN3, and IGHV1-24;IGHJ6-1, IGHV1-58;IGHJ3-1 and IGHV2-5;IGHJ4-1 for SP2 (**Fig. 5A**)^20, 21^. While these were the germlines predominant for neutralization, we aimed to interrogate the B cell repertoire to understand if the same or different germlines were employed for the Fc-functions evaluated in this study. The three most abundant germlines able to induce ADCP and ADCD against SARS-CoV-2 Wuhan were evaluated. Germlines showing same frequencies were included in the analyses. In SN2 cohort, we found the IGHV3-53;IGHJ6-1 (n=3; 13.0%), IGHV1-69;IGHJ3-1 (n=2; 8.7%), IGHV1-69;IGHJ4-1 (n=2; 8.7%), IGHV3-30;IGHJ4-1 (n=2; 8.7%), and IGHV3-30;IGHJ6-1 (n=2; 8.7%) (**Fig. 5B-C; Extended data Table 3**). These germlines accounted for 23.9% of the whole repertoire but the number of sequences recovered was too limited for a proper evaluation of this group. In the SN3 cohort, the most abundant germlines were the IGHV1- 46;IGHJ6-1 (n=20; 12.7%), IGHV1-69;IGHJ4-1 (n=17; 10.8%) and IGHV1-58;IGHJ3-1 (n=12; 7.6%), which constituted over 27.8% of the repertoire (**Fig. 5B-C; Extended data Table 3**). The SP2 cohort showed to predominantly use the IGHV1-69;IGHJ4-1 (n=15; 10.0%), IGHV1-69;IGHJ3-1 (n=8; 5.3%), IGHV2-5;IGHJ4-1 (n=8; 5.3%), IGHV1-46;IGHJ4-1 (n=6; 4.0%) and IGHV1-58;IGHJ3-1 (n=6; 4.0%) germlines which represented the 20.7% of the entire repertoire (**Fig. 5B-C; Extended data Table 3**). Interestingly, IGHV1-69;IGHJ4-1 was the only germline publicly shared across all three cohorts. In addition, SN3 and SP2 shared also the IGHV1-58;IGHJ3-1 germline, known to encode for broadly neutralizing antibodies^26^. Interestingly, around 60-70% of predominant germlines encoding for Fc-functions mediating antibodies are different from those used by neutralizing antibodies. Indeed, only one out of three germlines (IGHV1-58;IGHJ3-1; 33.3%) in the SN3 cohort and two out of five germlines (IGHV1-58;IGHJ3-1 and IGHV2-5;IGHJ4-1; 40.0%) in the SP2 group responsible for ADCP and ADCD were shared with germlines predominantly used to encode for neutralizing antibodies. Following, we compared the ability of predominant germlines to induce neutralization, ADCP and ADCD to Wuhan, Omicron BA.1 and BA.2 (**Fig. 5D-F**). The data confirmed that predominant germlines mediating Fc-functions are different from those involved in neutralization. All predominant germlines identified in the SN2 cohort showed low activities against tested variants (**Fig. 5D**). In the SN3 cohort, the IGHV1-46;IGHJ6-1 showed the strongest Fc- function activities against Wuhan and BA.1 variants. Interestingly, despite IGHV1-46;IGHJ6-1 antibodies show high ADCP and ADCD against BA.1, no neutralization was observed against this variant (**Fig. 5E**). Finally for the SP2 group, in line with the neutralization data, the most active germline was the IGHV2-5;IGHJ4-1 which showed high ADCP and ADCD against Wuhan and the BA.2 variant (**Fig. 5F**).

**Figure 5.**
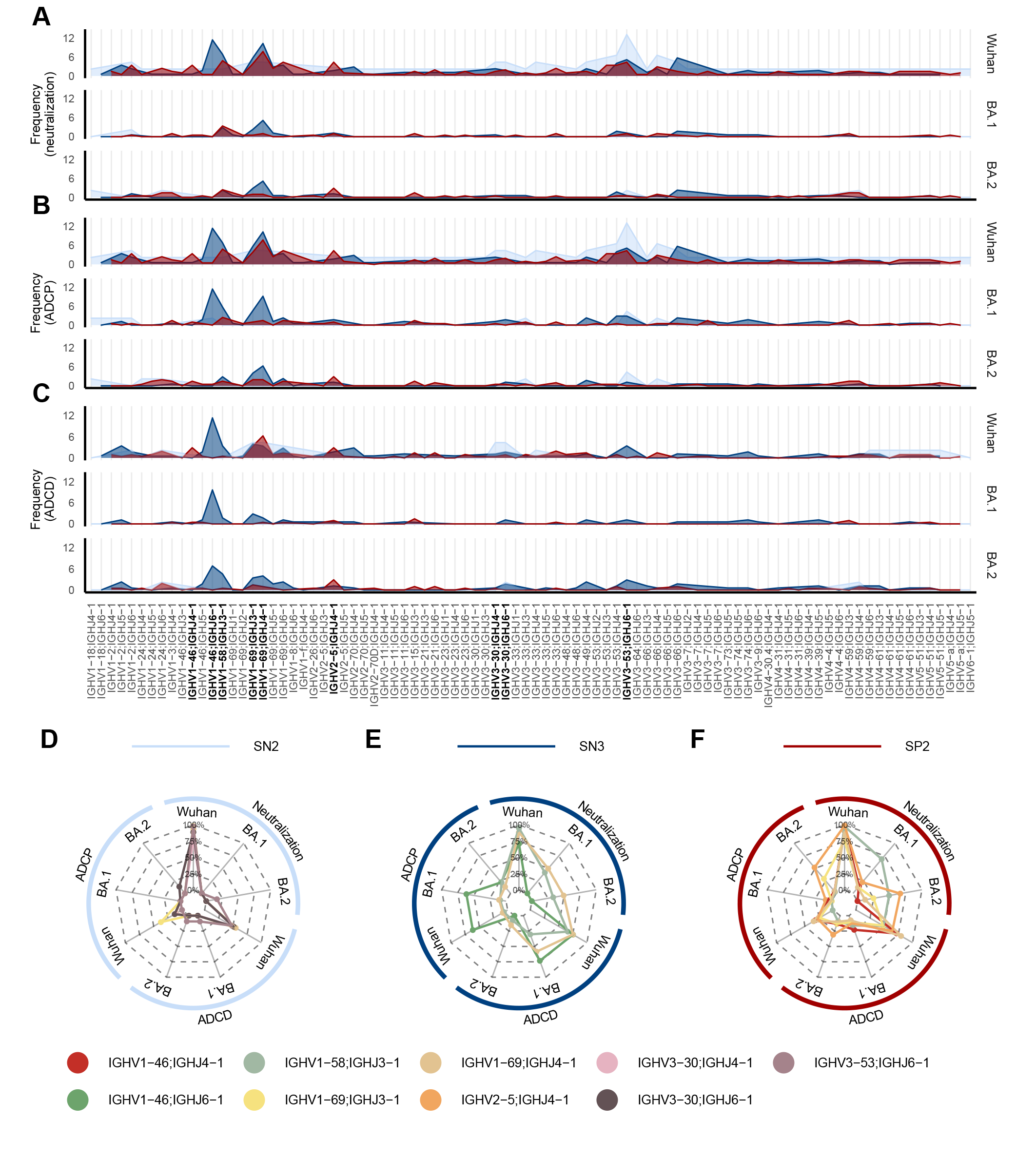
IGHV;IGHJ gene rearrangements implicated in ADCP and ADCD. (A-C) Graphs show the paired IGHV;IGHJ frequency usage of mAbs neutralization (A), ADCP (B) and ADCD (C) against the Wuhan, BA.1 and BA.2 viruses. Light blue, dark blue and red represent SN2, SN3 and SP2 cohorts respectively. In bold were highlighted the most abundant rearrangements used by antibodies able to drive ADCP and ADCD. (D-F) Radar plots describe the neutralization, ADCP and ADCD activities of predominant germlines identified in the SN2 (IGHV3-53;IGHJ6-1, IGHV1-69;IGHJ3-1, IGHV1-69;IGHJ4-1, IGHV3-30;IGHJ4-1 and IGHV3-30;IGHJ6-1) (D), SN3 (IGHV1-46;IGHJ6-1, IGHV1-69;IGHJ4-1 and IGHV1-58;IGHJ3-1) (E) and SP2 (IGHV1-69;IGHJ4-1, IGHV1-69;IGHJ3-1, IGHV2-5;IGHJ4-1, IGHV1-46;IGHJ4-1 and IGHV1-58;IGHJ3-1) (F). The percentages of functionality for neutralization, ADCP and ADCD are reported within each radar plot.

## DISCUSSION

Given the emerging evidence of the role of Fc-functions in protecting from COVID-19, we used a unique panel of 482 SARS-CoV-2 neutralizing human monoclonal antibodies to understand whether S protein domains and epitopes involved in Fc-functions differ from those mediating neutralization. With our panel we confirmed previous observations that antibodies that lose neutralization activity against variants can retain Fc-functions. In addition, we made the novel observation that Fc-functions are driven mostly by NTD and RBD Class 3- targeting nAbs while neutralization is known to be mediated mostly by RBD Class 1/2 antibodies. Differences between neutralization and Fc immune responses were also confirmed by our repertoire analyses showing the use of different B cell germlines for these two functions. Interestingly, despite SN3 and SP2 show similar levels of ADCP and ADCD and to target the same epitope and domain on the S protein, they employ different germlines. This observation highlights once more how the antigenic imprinting induced by vaccination or infection shapes differently the B cell and antibody response to SARS-CoV-2. Discrepancies between the neutralization and Fc-functions can also be observed by comparing the polyclonal (**Fig. 1**) and monoclonal responses (**Fig. 2A and 3A**) where the higher ADCP and ADCD activities of the SP2 polyclonal response is not reflected at monoclonal level. This suggests that in the polyclonal response not-neutralizing antibodies play an important role in driving Fc-functions. In conclusion, our study unravels key features of the Fc-dependent antibody effector functions and highlight the intersection between neutralization and Fc-mediated activities in subjects after vaccination or with hybrid immunity. The high-resolution picture provided in this study strongly suggests that Fc-mediating domains and epitopes should be considered to define novel correlates of protection, design of improved vaccines and selection of antibodies against COVID-19.

## METHODS

### Enrollment of COVID-19 vaccinees and human sample collection

Thanks to a collaboration with the Azienda Ospedaliera Universitaria Senese, Siena (IT), human samples were collected from healthy donors immunized with two or three vaccine doses and from COVID-19 convalescent who had been infected before received two vaccine doses. All subjects were immunized with the BNT162b2 mRNA vaccine, except for one subject which received the mRNA-1273 vaccine as third dose. All the participants gave their written consent and both sexes were included in this study. This work was approved by the Comitato Etico di Area Vasta Sud Est (CEAVSE) ethics committees (Parere 17065 in Siena) and performed according to good clinical practice and in line with the declaration of Helsinki (European Council 2001, US Code of Federal Regulations, ICH 1997). The study was unblinded and not randomized. No statistical methods were used to predetermine sample size.

### Single-cell RT-PCR and Ig gene amplification

SARS CoV-2 neutralizing mAbs were expressed as previously described^21^. Briefly, reverse transcription polymerase chain reaction (RT-PCR) was performed using the cell lysate derived from the 384-cell sorting plate. Then, PCRI and nested PCRII were performed. PCRII products were purified and used to perform Gibson Assembly ligation in human Igγ1, Igκ, and Igλ expression vectors as previously described^27^. Next, transcriptionally active PCR (TAP) was performed using of Q5 High-Fidelity DNA Polymerase. TAP products were then purified and used for transient transfection in the Expi293F cell line following manufacturing instructions.

### Antibody-dependent cellular phagocytosis (ADCP) assay

Flow cytometry-based assay was used to analyze Antibody-dependent cellular phagocytosis (ADCP). Briefly, 200 µg of stabilized histidine tagged S protein were labelled using Strep-Tactin™XT Conjugate DY-649 (IBA Lifesciences) and coated with 1 mg of magnetic beads (Dynabeads His-Tag, Invitrogen) as previously described^21^. The S protein-beads were incubated with nAbs or heated-inactivated plasma diluted in complete RPMI (1:40) for 1 hour at room temperature. Then, S protein-beads were incubated with the monocytic THP-1 cell line overnight, fixed fixation buffer (Biolegend) and acquired on BD LSR II flow cytometer (Becton Dickinson). Single technical replicates were performed for each experiments. Plasma from a pre-pandemic healthy subject was used as negative control and phagocytosis score was calculated as the percentage of THP-1 cells that engulfed fluorescent beads multiplied by the median fluorescence intensity of the population.

### Antibody-dependent Complement Deposition (ADCD) assay

To investigate the antibodies ability to induce complement deposition, Expi293F cells (Thermo Fisher, Cat#A14527) were transiently transfected with SARS-CoV-2 original S protein, BA.1 or BA.2 expression vectors (pcDNA3.1_ spike_del19) using Expifectamine Enhancer according to the manufacturer’s protocol (Thermo Fisher). Two days later, heated-inactivated plasma diluted 1/50 in complete Expi medium (Thermo Fisher) or monoclonal antibodies were incubated with HEK-S protein cells for 30 minutes at 37°C, with 5% CO_2_ and 120 rpm shaker speed. Then, 50 µl of Expi medium containing 6% of baby rabbit complement (Cederlane) was added and cells were incubated at 37°C, with 5% CO_2_ and 120 rpm shaker speed for 30 minutes. After incubation, cells were washed with PBS and stained with goat anti-rabbit polyclonal antibody against C3 (MP Biomedicals) for 1h on ice. Cells were fixed with fixation buffer (Biolegend) for 15 minutes on ice. Then, cells were washed, resuspended in 100 µL of PBS1X and acquired using BD LSR II flow cytometer (Becton Dickinson). Plasma isolated from a pre-pandemic healthy subject was used as negative control. Single technical replicates were performed for each experiments. Results were reported as median fluorescence intensity of C3 deposition on the cells.

### Functional repertoire analyses

nAbs VH and VL sequence were manually curated and retrieved using CLC sequence viewer (Qiagen). Aberrant sequences were eliminated from the data set. Then, analyzed reads were saved in FASTA format and Cloanalyst software was used to perform the repertoire analyses (http://www.bu.edu/computationalimmunology/research/software/). The figure was assembled with ggplot2 v3.3.5.

### Statistical analysis

GraphPad Prism Version 8.0.2 (GraphPad Software, Inc., San Diego, CA) was used to perform Statistical analysis. To evaluate statistical significance between the cohorts examined in this study a Nonparametric Mann-Whitney t test was applied. Statistical significance was indicated as * for values ≤ 0.05, ** for values ≤ 0.01, and *** for values ≤ 0.001.

### Data availability

Source data are provided with this paper. All data supporting the findings in this study are available within the article or can be obtained from the corresponding author upon request. SARS-CoV-2 antibody sequences were deposited and accessible from https://github.com/dasch-lab/SARS-CoV-2_nAb_third_dose.

## Acknowledgments

This work received funding by the European Research Council (ERC) advanced grant (agreement number 787552 (vAMRes)), and the Italian Ministry of Health (COVID-2020-12371817 project). In addition, this work was supported by a fundraising activity promoted by Unicoop Firenze, Coop Alleanza 3.0, Unicoop Tirreno, Coop Centro Italia, Coop Reno e Coop Amiatina.

## Author contributions

Conceived the study: I.P., E.A. and R.R.; Performed ADCP and ADCD assays and analyzed data: I.P.; Expression and production of S proteins: E.P; Performed repertoire analyses: E.A. and G.M.; Manuscript writing: I.P., E.A. and R.R.; Final revision of the manuscript: I.P., G.M., E.P., E.A. and R.R.; Coordinated the project: E.A. and R.R.

## Additional information

Correspondence and requests for materials should be addressed to E.A. and R.R.

## EXTENDED DATA FIGURES

**Extended Data Figure 1.**
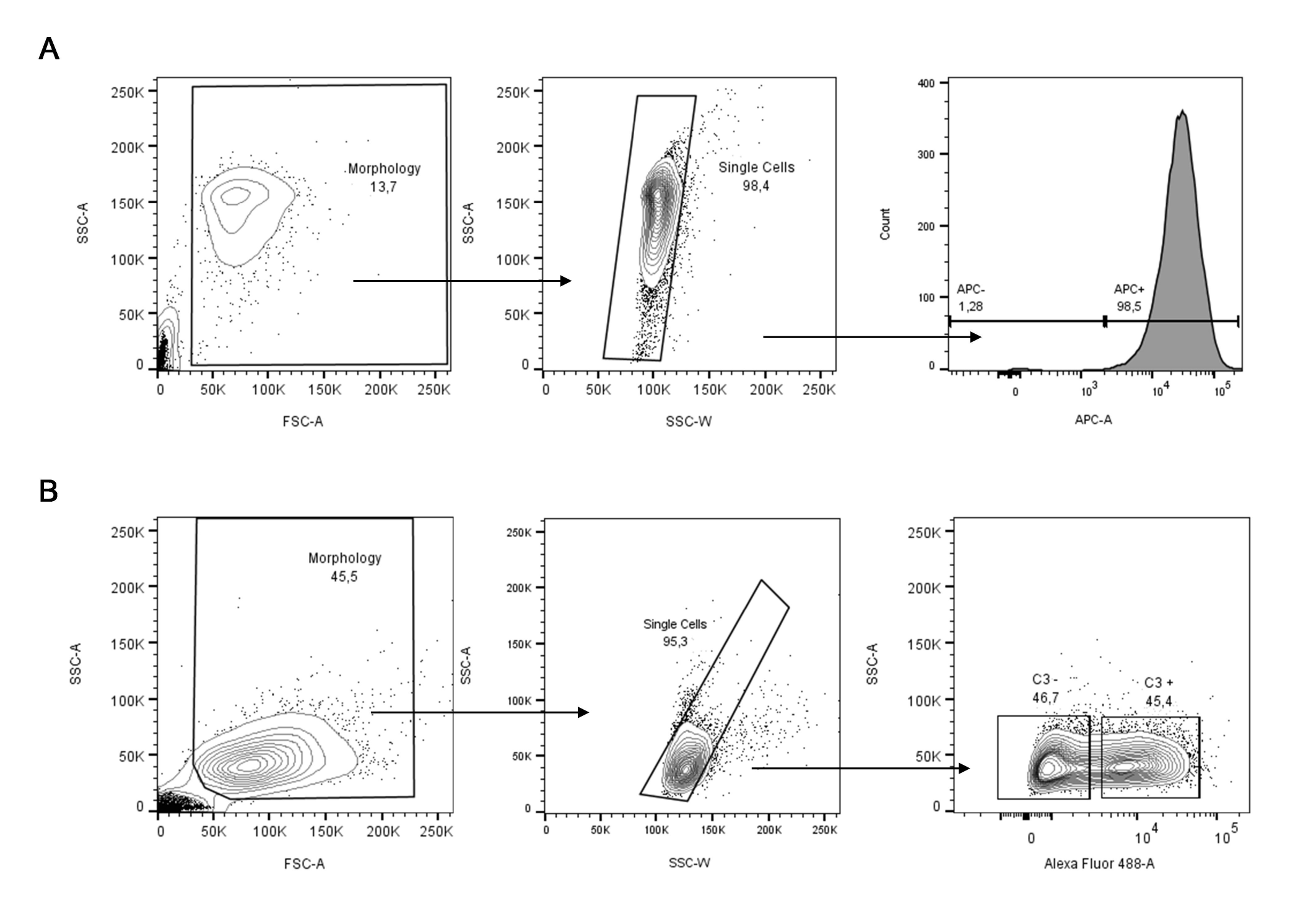
Gating strategy for ADCP and ADCD. (A) The gating strategy used to detect fluorescent THP-1 cells is composed by: Morphology; Single cells; APC^+^ cells. (B) Description of successive subpopulations of cells selected for ADCD analysis in flow cytometry is composed by morphology, Single cells, A488^+^ subsets.

**Extended Data Figure 2.**
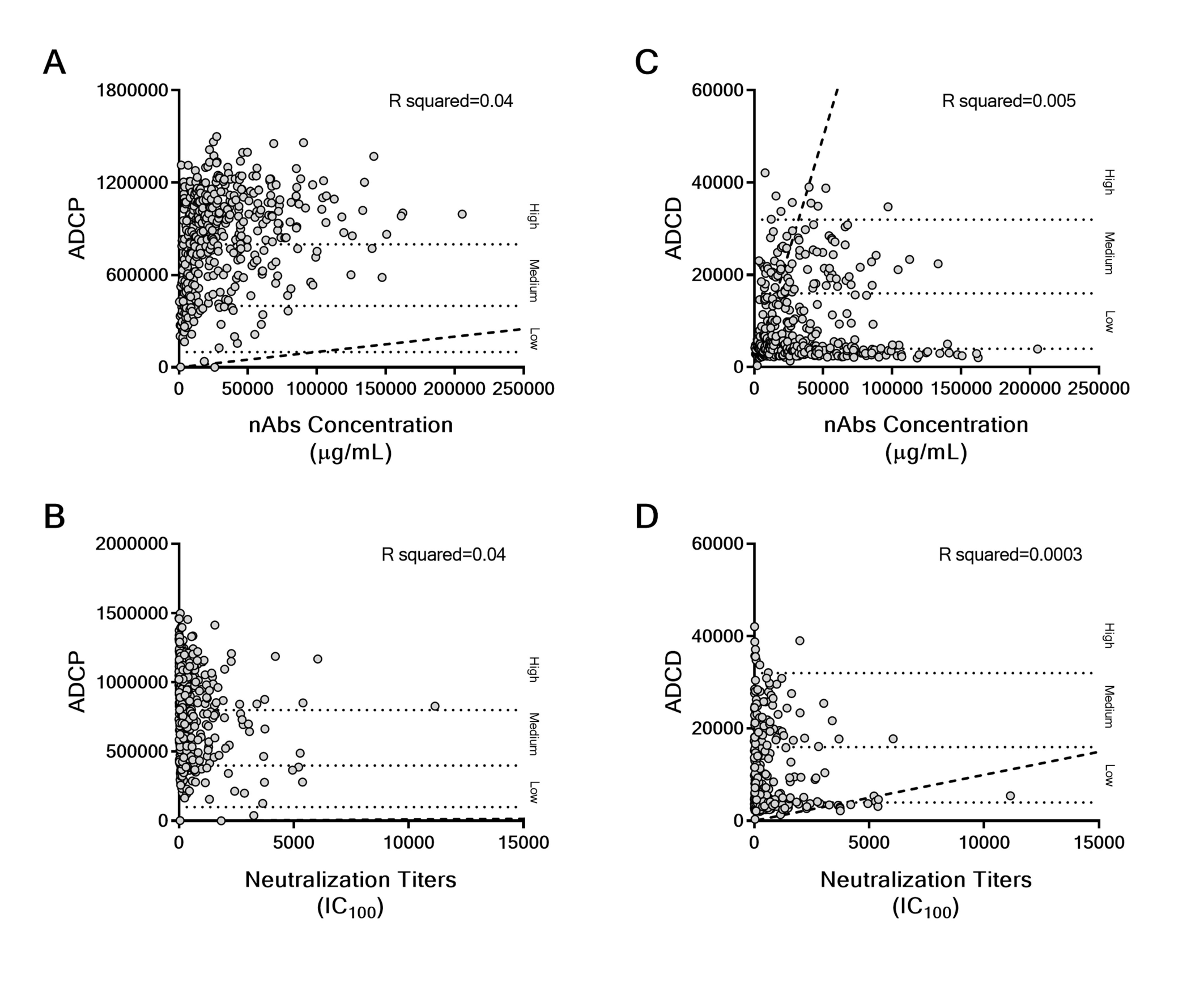
Correlation between Fc-function nAbs concentration and neutralization titers. (A) Concentration-dependent effect of nAbs on ADCP. (B) Correlation between ADCP and neutralization. (C) Concentration-dependent effect of nAbs on ADCD. (D) Correlation between ADCD and neutralization. Statistical analyses were performed used Pearson correlation coefficient and a linear correlation is observed if the R- squared value ranges from 0 to 1.

## SUPPLEMENTARY TABLES

**Extended Data Table 1.**
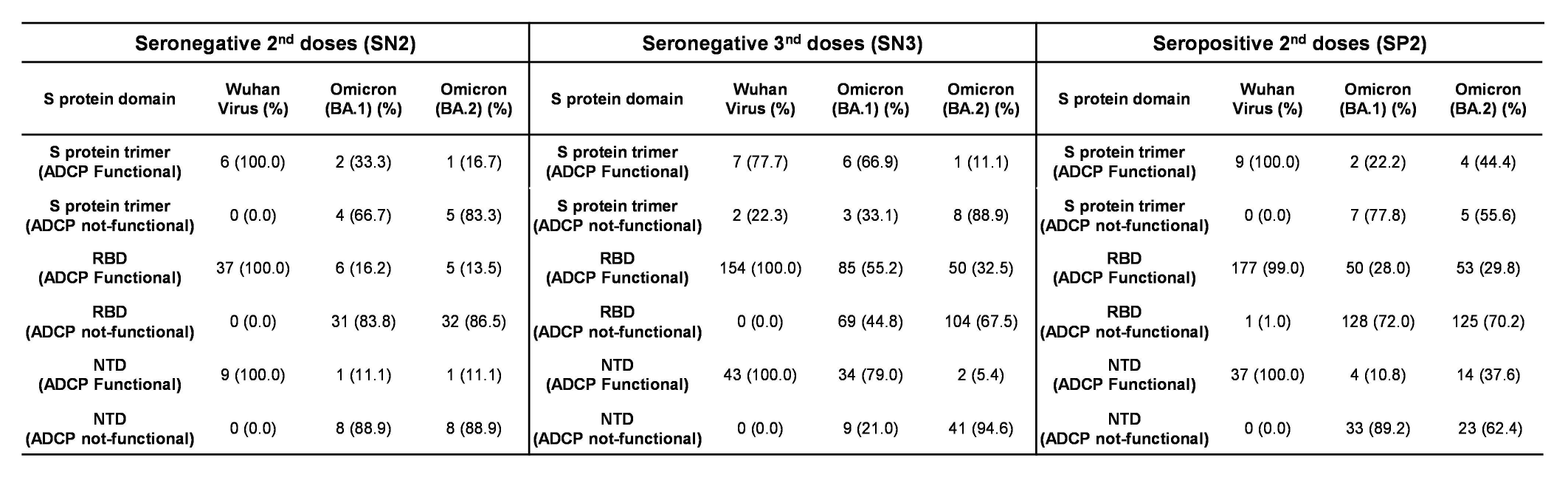
Summary of ADCP functional and not-functional nAbs. The table summarizes the number and percentages of S protein trimer, RBD and NTD targeting nAbs phagocytosis inducers against the original Wuhan virus, the Omicron variants BA.1 and BA.2 in SN2, SN3 and SP2 cohort.

**Extended Data Table 2.**
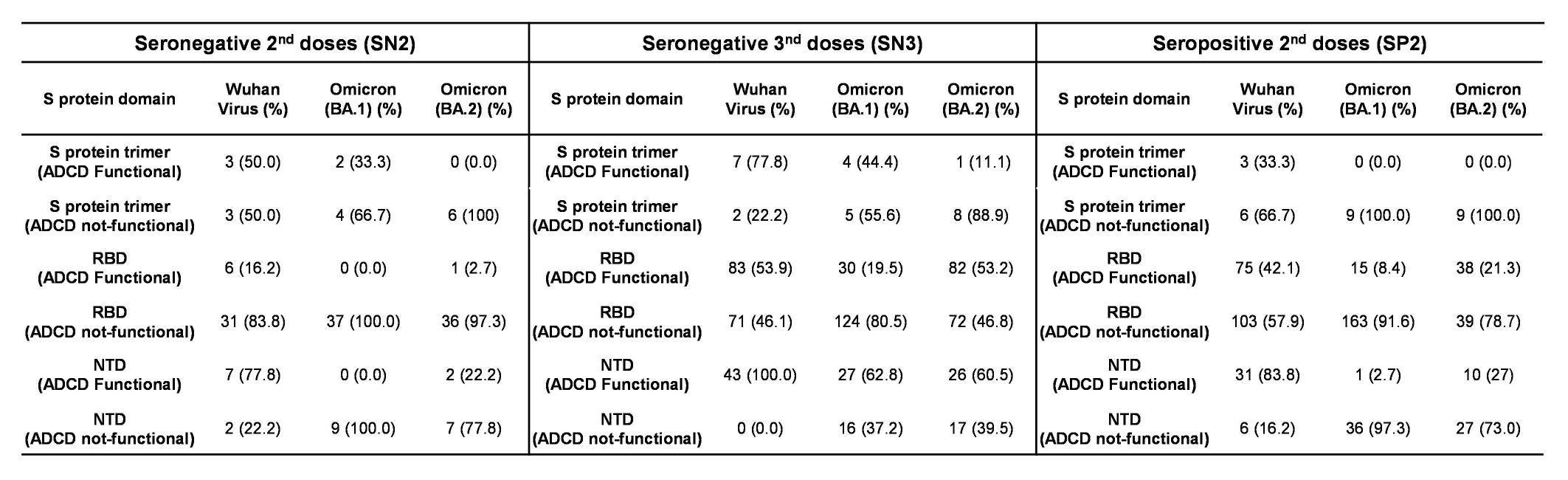
Summary of ADCD functional and not-functional nAbs. The table summarize the number and percentages of nAbs S protein trimer, RBD and NTD binders able to induce complement deposition against the SARS-CoV-2 original virus, the Omicron variants BA.1 and BA.2 in SN2, SN3 and SP2 cohort.

**Extended Data Table 3.** Predominant germlines involved in ADCP and ADCD. The table show the number and frequency of the predominant IGHV;IGHJ rearrangement involved in ADCP and ADCD against the original Wuhan virus.

| Seronegative 2<sup>nd</sup> doses (SN2) |  |  |
| --- | --- | --- |
| IGHV;IGHJ<br>Rearrangement | Number | Frequency |
| IGHV3-53;IGHJ6-1 | 3 | 13.40 |
| IGHV1-69;IGHJ3-1 | 2 | 8.70 |
| IGHV1-69;IGHJ4-1 | 2 | 8.70 |
| IGHV3-30;IGHJ4-1 | 2 | 8.70 |
| IGHV3-30;IGHJ6-1 | 2 | 8.70 |
| Seronegative 3<sup>nd</sup> doses (SN3) |  |  |
| IGHV;IGHJ<br>Rearrangement | Number | Frequency |
| IGHV1-46;IGHJ6-1 | 20 | 12.66 |
| IGHV1-69;IGHJ4-1 | 17 | 10.76 |
| IGHV1-58;IGHJ3-1 | 12 | 7.59 |
| Seropositive 2<sup>nd</sup> doses (SP2) |  |  |
| IGHV;IGHJ<br>Rearrangement | Number | Frequency |
| IGHV1-69;IGHJ4-1 | 15 | 10.00 |
| IGHV1-69;IGHJ3-1 | 8 | 5.33 |
| IGHV2-5;IGHJ4-1 | 8 | 5.33 |
| IGHV1-46;IGHJ4-1 | 6 | 4.00 |
| IGHV1-58;IGHJ3-1 | 6 | 4.00 |

## REFERENCES

1. WHO. Coronavirus (COVID-19) Dashboard. (2023).

2. Jeong, H.W., et al. Enhanced antibody responses in fully vaccinated individuals against pan-SARS-CoV-2 variants following Omicron breakthrough infection. Cell reports. Medicine 3, 100764 (2022).

3. Arora, P., et al. Omicron sublineage BQ.1.1 resistance to monoclonal antibodies. The Lancet Infectious Diseases 23, 22–23 (2023).

4. Khoury, D.S., et al. Neutralizing antibody levels are highly predictive of immune protection from symptomatic SARS-CoV-2 infection. Nature Medicine 27, 1205–1211 (2021).

5. Pušnik, J., et al. SARS-CoV-2 humoral and cellular immunity following different combinations of vaccination and breakthrough infection. Nature Communications 14, 572 (2023).

6. Zhou, H., et al. Sensitivity to Vaccines, Therapeutic Antibodies, and Viral Entry Inhibitors and Advances To Counter the SARS-CoV-2 Omicron Variant. Clinical Microbiology Reviews 35, e00014–00022 (2022).

7. Gruell, H., et al. SARS-CoV-2 Omicron sublineages exhibit distinct antibody escape patterns. Cell host & microbe 30, 1231–1241.e1236 (2022).

8. Nilles, E.J., et al. Tracking immune correlates of protection for emerging SARS-CoV-2 variants. The Lancet Infectious Diseases 23, 153–154 (2023).

9. Sigal, A. Milder disease with Omicron: is it the virus or the pre-existing immunity? Nature reviews. Immunology 22, 69–71 (2022).

10. Mackin, S.R., et al. Fc-γR-dependent antibody effector functions are required for vaccine-mediated protection against antigen-shifted variants of SARS-CoV-2. Nature microbiology 8, 569–580 (2023).

11. Zhang, A., et al. Beyond neutralization: Fc-dependent antibody effector functions in SARS-CoV-2 infection. Nature Reviews Immunology 23, 381–396 (2023).

12. Amin, A., et al. Therapeutic and vaccine-induced cross-reactive antibodies with effector function against emerging Omicron variants. bioRxiv, 2023.2001.2017.523798 (2023).

13. Mackin, S.R., et al. Fc-γR-dependent antibody effector functions are required for vaccine-mediated protection against antigen-shifted variants of SARS-CoV-2. Nature microbiology 8, 569–580 (2023).

14. Pušnik, J., et al. SARS-CoV-2 humoral and cellular immunity following different combinations of vaccination and breakthrough infection. Nat Commun 14, 572 (2023).

15. Bowman, K.A., et al. Hybrid Immunity Shifts the Fc-Effector Quality of SARS-CoV-2 mRNA Vaccine- Induced Immunity. mBio 13, e0164722 (2022).

16. Jegaskanda, S., Vanderven, H.A., Wheatley, A.K. & Kent, S.J. Fc or not Fc; that is the question: Antibody Fc-receptor interactions are key to universal influenza vaccine design. Human Vaccines & Immunotherapeutics 13, 1288–1296 (2017).

17. DiLillo, D.J., Palese, P., Wilson, P.C. & Ravetch, J.V. Broadly neutralizing anti-influenza antibodies require Fc receptor engagement for in vivo protection. The Journal of clinical investigation 126, 605–610 (2016).

18. Gunn, B.M., et al. A Role for Fc Function in Therapeutic Monoclonal Antibody-Mediated Protection against Ebola Virus. Cell host & microbe 24, 221–233.e225 (2018).

19. Troisi, M., et al. A new dawn for monoclonal antibodies against antimicrobial resistant bacteria. Frontiers in microbiology 13, 1080059 (2022).

20. Andreano, E., et al. mRNA vaccines and hybrid immunity use different B cell germlines against Omicron BA.4 and BA.5. Nature Communications 14, 1734 (2023).

21. Andreano, E., et al. Hybrid immunity improves B cells and antibodies against SARS-CoV-2 variants. Nature 600, 530–535 (2021).

22. Andreano, E., et al. B cell analyses after SARS-CoV-2 mRNA third vaccination reveals a hybrid immunity like antibody response. Nature Communications 14, 53 (2023).

23. Torres, J.L., et al. Structural insights of a highly potent pan-neutralizing SARS-CoV-2 human monoclonal antibody. Proceedings of the National Academy of Sciences 119, e2120976119 (2022).

24. Pinto, D., et al. Cross-neutralization of SARS-CoV-2 by a human monoclonal SARS-CoV antibody. Nature 583, 290–295 (2020).

25. Yuan, M., et al. A highly conserved cryptic epitope in the receptor binding domains of SARS-CoV-2 and SARS-CoV. Science 368, 630–633 (2020).

26. Zhou, T., et al. Structural basis for potent antibody neutralization of SARS-CoV-2 variants including B.1.1.529. Science 376, eabn8897 (2022).

27. Andreano, E., et al. Extremely potent human monoclonal antibodies from COVID-19 convalescent patients. Cell 184, 1821–1835.e1816 (2021).

